# Mind Your Spectra: Points to be Aware of when Validating the Identification of Isobaric Histone Peptidoforms

**DOI:** 10.1101/2024.11.13.623238

**Authors:** Hassan M. Hijazi, Julie Manessier, Sabine Brugière, Tina Ravnsborg, Marie Courçon, Baptiste Brulé, Karine Merienne, Ole N Jensen, Anne-Marie Hesse, Christophe Bruley, Delphine Pflieger

## Abstract

Mass spectrometry (MS) has become a central technique to identify and quantify post-translational modifications (PTMs), overcoming limitations of antibody-based methods. Histones get dynamically modified by diverse chemical groups, particularly on their numerous lysine residues, to fine-tune all DNA-templated processes. Reliable identification of histone PTMs remains challenging and still requires manual data curation. This study focused on the Lys27-Arg40 stretch of histone H3, considered four sequence variants, an increasing number of lysine PTMs and artifacts coming from histone sample processing, which resulted in many peptides with the same atomic composition. Our analysis revealed the value of low-mass b1 and cyclic immonium fragment ions to validate identification of the distinct peptidoforms. We examined how MS/MS spectra are transformed by common software tools during the conversion of RAW files into peak lists, and highlighted how some parameters may erase the informative low-mass fragments. We established the fragmentation profiles and retention times for forty H3 K27-R40 variant×PTM combinations, including the mouse-specific variants H3mm7 and H3mm13, and targeted their detection in histone samples extracted from mouse testis and brain via a scheduled parallel reaction monitoring (PRM) analysis. The transcripts of these two mouse-specific variants were reported to be highly abundant in these tissues and the corresponding proteins may seem to be identified by data-dependent analyses. However, we only detected very low levels of the unmodified form of H3mm7 and found no trace of H3mm13 by PRM. Our work contributes to reliably deciphering the histone code shaped by distinct sequence variants and numerous combinations of PTMs.

## Introduction

Liquid chromatography coupled to tandem mass spectrometry (LC-MS) has become the method of choice for high-throughput analysis of post-translational modifications (PTMs) (Huang et al., 2015). The complexity of PTM analysis is particularly evident in histone proteins because they are extensively modified by a multitude of PTMs, especially on lysine residues. These residues can be modified by acetylation, methylation, ubiquitination, and a range of acylations of variable lengths (Nitsch et al., 2021). Histones are particularly rich in lysine (K) and arginine (R) residues and are consequently usually submitted to a derivatization step before trypsin digestion, to avoid their extensive cleavage into very small peptides (Garcia et al., 2007; Maile et al., 2015; Meert et al., 2016). Propionic anhydride is widely used to add a propionyl group to unmodified and monomethylated lysines, forcing trypsin to follow the Arg-C cleavage rule and produce longer, consistent, and more hydrophobic peptides that are more amenable to LC-MS analysis (Garcia et al., 2007). However, this reaction requires a controlled pH of *∼* 8, otherwise, propionylation at the desired sites may fail (underpropionylation) or excessive off-target propionylation (overpropionylation) may be induced on serine (S), threonine (T) and tyrosine (Y) residues (Garcia et al., 2007; Sidoli and Garcia, 2017; Meert et al., 2015; Searfoss et al., 2023). During data analysis, the large number of PTMs combined with subtle sequence variations and propionylation artifacts populate the search space used to interpret fragmentation spectra with many strictly isobaric peptides derived from histone peptides. This makes validating the peptide identifications suggested by software tools very challenging.

Conventional shotgun proteomics relies on a targetdecoy strategy to control the false discovery rate and thus distinguish correct from incorrect peptide-spectrum matches. However, there exists no proper equivalent for modified peptides produced from a few selected protein sequences. Neither specifying cut-off identification scores nor a PTM localization probability threshold proves highly efficient to filter false identifications. As of today, several strategies have been devised to tackle various aspects of the limitations associated with the identification and quantification of histone PTMs. For example, some efforts have attempted to alleviate potential ambiguities in database search results obtained on histones. One approach modeled all possible variant×PTM combinations (i.e., peptidoforms) from a histone sequence database, considering only known-toexist PTMs and filtering out those not matching the mass-to-charge ratios found in peak lists extracted from RAW files. Subsequently, a few sets of PTMs were prioritized for use in parallel database searches to maximize peptide identifications, particularly for peptides with co-occurring PTMs (Willems et al., 2017). Another study used a histone deacetylase (HDAC) to catalytically remove acetyl groups from histone lysines, and empirically flag false identifications by quantifying the enzyme-induced changes of PTMs over time. With this strategy, it was possible to spot incorrect annotations that did not respect the expected decline in acetylated peptides versus the rise in unmodified peptides in response to the enzyme’s activity (De Clerck et al., 2021). Finally, the histone-tailored software, Epiprofile, was specifically developed to identify and quantify a set of modified histone peptides including positional isomers based on prior knowledge of the order of elution of PTMs and unique discriminating fragment ions (Yuan et al., 2015, 2018). The quality of the data determines how well Epiprofile performs, and adding PTMs that are not covered by the built-in elution order rule requires programming expertise (Thomas et al., 2020). Overall, several strategies have been devised to tackle limitations associated with the reliability of identification and quantification of histone PTMs.

In the present study, we sought to define robust rules to allow the correct identification of histone peptide sequences by LC-MS/MS when considering an extended search space due to subtle sequence variations combined with several lysine acylations. Mouse-specific variants of histone H3 have been described, among which H3mm7 and H3mm13 were shown to be of high abundance at the transcript level in different organs, including mouse brain and testis (Maehara et al., 2015); in parallel, H3mm18 and H3mm7 have been attributed specific functions in skeletal muscle (Harada et al., 2018; Hirai et al., 2022). A first step to testing the biological relevance of such variants consists of assessing their abundance at protein level compared to canonical H3 and variant H3.3. We selected H3mm7 and H3mm13, and specifically analyzed the K27-R40 stretch, which differs by one or two amino acids across the four histone variants. We emphasize the importance of performing reversepropionylation of S/T residues because otherwise, excessive levels of derivatization lead to a dramatic increase in the number of erroneous identifications. We underscore the importance of detecting low-mass fragment ions in MS/MS spectra, particularly the b1 fragment and the CycIm ion, indicative of acyl and monomethyl modifications on H3K27, to increase confidence in peptide identity. Finally, using PRM analysis, we targeted forty K27-R40 peptidoforms of H3 histones extracted from mouse testis and brain, to definitively ascertain the presence of H3mm7 and H3mm13 variants.

## Materials and Methods

### Tissues and Synthetic Peptides

Brain and testis tissues were obtained from 2.5 to 3-month-old male mice (C57BL/6; see Supplementary Data Section 1). Synthetic peptides were purchased from JPT Peptide Technologies (Germany) and Synpeptide (China). The indexed retention time (iRT) kit was purchased from Biognosys (Switzerland).

### Acid Extraction of Histones

Brain hemispheres were kept intact, whereas testes were ground on dry ice using a pestle and mortar before histone extraction. Both tissues were resuspended in 0.2 M sulfuric acid (H_2_SO_4_) and sonicated using a 3-mm probe CV18 sonicator for 12 cycles of 5 s ON/5 s OFF at 20% amplitude. After incubating the lysed cells for 90 min, the supernatant containing dissolved histones was precipitated on ice for 30 min after adding trichloroacetic acid (TCA) to a final concentration of 20%. Precipitated histones were centrifuged (15 min, 18,400 *g*, 4 °C) and subsequently washed twice without resuspending the precipitate: once with ™20 °C 0.1% hydrochloric acid (HCl) in acetone and then with ™20 °C pure acetone. Finally, the samples were dried under the hood for 15 min and resuspended in 160 µL SDS-PAGE loading buffer. A 10 ™12 µL aliquot was loaded onto a 4-12% polyacrylamide gel and submitted to electrophoresis. Bands corresponding to H3 were excised under the hood and cut into three 1-mm cubes using a scalpel before storing at ™20 °C.

### In Vitro Propionylation and Tryptic Digestion of Histones

We largely followed the protocol described by Maile et al (Maile et al., 2015) with slight modifications (Geshkovski et al., 2024). Briefly, propionic anhydride solution was freshly prepared before each derivatization step by diluting pure propionic anhydride 1:100 in water then diluting 100 mL of this in 1 mL of 100 mM TEAB pH 8.5 (for the synthetic histone peptides, this solution was immediately applied). Gel pieces containing trapped histone proteins were subjected to two rounds of derivatization by incubating in 100 µL of derivatization reagent with agitation for 30 min. Derivatized histones were subsequently digested overnight at 37 °C with trypsin (V5111 from Promega) diluted in 50 mM TEAB pH 8.5 and added at an estimated enzyme:substrate ratio of 1:20. A further two rounds of derivatization were performed to cap the newly generated histone peptide N-termini. Reversepropionylation to remove undesired propionylation on S/T/Y residues was conducted by applying 20 µL of 500 mM hydroxylamine and 12 µL of ammonium hydroxide (pH 12) for 20 min at room temperature under agitation. The reaction was stopped by adding a few drops of pure trifluoroacetic acid (TFA). Samples were vacuum-dried before desalting using Affinisep SPE tips (BioSPE^®^ PurePep). The prepared peptides were stored at ™20 °C until MS analysis.

### Liquid Chromatography-Mass Spectrometry Acquisition

For LC-MS analysis, histone peptides were reconstituted in LC buffer (0.1% v/v TFA, 5% v/v acetonitrile [ACN] in water) before loading onto a PepMap C18 precolumn (300 µm × 5 mm, Dionex, Thermo Fisher Scientific) with 0.1% formic acid in 2 min. Histone peptides were then separated on a C18 reverse-phase capillary column (Aurora C18, 75 µm×25 cm, 1.7 µm beads, 120 Å pore size) in an Ultimate 3000 nano chromatography system (Thermo Fisher Scientific) coupled to a QExactive HF mass spectrometer (Thermo Fisher Scientific). The capillary column flow rate was set to 300 nL/min. The mobile phases consisted of solvent A (water with 0.1% formic acid) and solvent B (ACN with 0.08% (v/v) formic acid). Peptides were separated by applying a gradient consisting of an increase in solvent B from 2% to 7% in 5 min, then from 7% to 31% over 55 min, from 31% to 41% over 8 min, followed by a 9-min flush of the column at 72.2% B and re-equilibration at 2% B for a total run time of 92 min.

For data-dependent acquisition (DDA), MS1 spectra (from m/z 300 to 1300) were acquired at a resolution of 60,000, an AGC target of 10 ^6^ and a maximum injection time (max IT) of 200 ms. MS/MS spectra were acquired at a resolution of 15,000, an AGC target value of 2 *×* 10 ^5^, with a max IT of 100 ms, a loop count of 20, and a 1.5-m/z isolation window. Precursor ions were fragmented using higher-energy collisional dissociation (HCD) at 30% normalized collision energy. The first fixed mass was set to 80 m/z and dynamic exclusion time to 10 s. For PRM analyses, the mass spectrometer automatically switched between one MS1 scan and 14 MS/MS scans. An inclusion list of the m/z values of interest along with the retention times (RT) of their respective modified histone peptide precursor ions was provided (See Supplementary Table S1). PRM MS/MS spectra were acquired at a resolution of 30,000, with an AGC target of 10 ^6^, a max IT of 120 ms, and an isolation window set to 1.6 m/z.

Middle-down analysis was performed on an Orbitrap Eclipse mass spectrometer coupled inline to a Neo Vanquish LC (Thermo Fisher Scientific). Histones were extracted from mouse testis as described above and digested overnight with Glu-C endoprotease (V1651, New England BioLabs) at 37 °C, at an enzyme:substrate ratio of 1:20, in 0.05 M ammonium acetate (pH 4). Samples were loaded onto a trap column (Thermo Fisher Scientific, PepMap Neo; 300 µm internal diameter, 5 mm length, C18 reverse-phase material, 5 µm diameter beads, and 100 Å pore size) and then eluted online onto an analytical hydrophilic interaction liquid chromatography (HILIC) column (Pulled Emitter, I.D. 75 µm) packed in-house with Polycat A (2 µm, 1000 Å, PolyLc inc.) to a length of 30 cm. The mobile phase consisted of solvent A (20 mM formic acid, 75% ACN, pH 4.5) and solvent B (1% formic acid in water). Peptides were separated by applying a gradient consisting of an increase in solvent B from 10% to 50% in 15 min, then from 50% to 65% in 70 min, and finally to 100% in 20 min. Data were acquired in the Orbitrap for both precursor and product ions, with a mass resolution of 120,000 for MS (max IT 100 ms, 500-1000 m/z scan range) and 30,000 for MS/MS (max IT 800 ms, 665-755 m/z scan range) to include only charge state 8+ of the H3 tail. The isolation window was set to 1.4 m/z. Electron Transfer Dissociation (ETD) reactions were carried out for 60-100 ms with ETD reagent target values of 40,000 and max IT of 1000 ms.

### Database Searches, Preprocessing, Quantification and Data Visualization

MS acquisition RAW files were converted to *mzdb* file format using mzdbwizard. To obtain the peak lists as mgf, the MGFBoost computing method (paper in preparation) was used within mzdbwizard. The MGF files were subsequently searched against an in-house curated histone (El Kennani et al., 2017) and contaminant databases using the Mascot search engine (Matrix Science v2.8). Enzyme specificity was set to Arg-C with no missed cleavage. N-terminal propionylation was set as a fixed modification whereas lysine acetyl (Kac), propionyl (Kpr), butyryl (standing for Kmonomethyl+propionyl; Kme1), di- and trimethyl (Kme2/me3), crotonyl (Kcr), hydroxybutyryl (Khb) and lactyl (Kla) were set as variable modifications. Mass tolerances for peptides and MS/MS fragments were set to 5 ppm and 10 ppm, respectively. For middle-down analyses, Mascot was used to search RAW files with the following settings: enzyme specificity set to Gluc-C (V8E), missed cleavage set to 1, and tolerances on precursor and fragment masses set to 1.03 Da and 20 ppm, respectively. The variable modifications excluded the derivatization-induced modifications and only included acetylation and methylations.

To compare the preprocessing algorithms, data were acquired on a mixture of synthetic peptides covering the stretch K27-R40 from the four H3 variants. Spectra in Figure 2 and 3 (see also Supplementary Figure S1) were extracted from their respective MGF files and annotated using custom scripts via *pyteomics* (Levitsky et al., 2018) and *spectrum_utils* (Bittremieux, 2019) python libraries, respectively.

**Figure 1.**
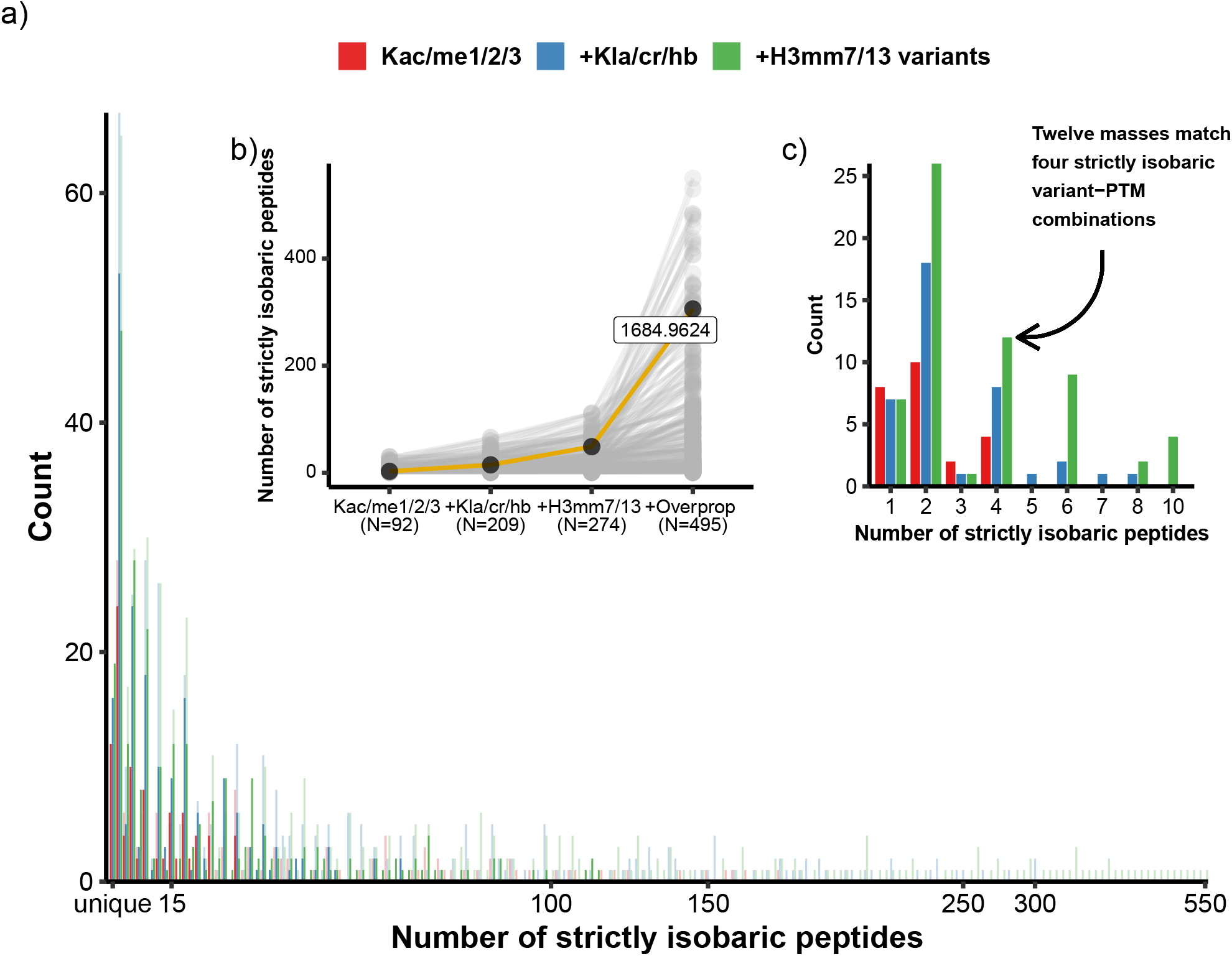
The number of strictly isobaric H3 K27-R40 masses increases considerably when novel PTMs, variants, and overpropionylation are allowed. (**a**) The number of isobaric peptides defined as a given (K27-R40 sequence x PTM combination on K27/K36/K37) was counted after successively adding variables: considering only classical PTMs (Kac and Kme1/2/3) on canonical H3 and variant H3.3 (Kac/me1/2/3; red bars); adding novel PTMs to H3 and H3.3 (+Kla/cr/hb; blue bars); considering two H3mm variants (+H3mm7/13 variants; green bars) and finally allowing for overpropionylation on S/T residues (same color code but translucent). This graph illustrates the H3 K27-R40 global search space from which a program filters precursor masses before assigning peptide-spectrum matches (PSMs). The values on the x-axis are not continuous as some are skipped when their count is zero. Inset (**b**) shows how the number of isobaric variant×PTM combinations increases, each line represents a unique mass. The mass 1684.96244 amu is highlighted, others are grayed out. The number of variant×PTM combinations strictly matching this mass increases when acylations (15 combinations), mouse-specific variants (49 combinations), and most prominently overpropionylation (306 combinations) are included. ‘N’ represents the total number of unique masses in each condition. Inset (**c**) shows a subsection of the “global search space” after adding some filters: (1) No more than one novel acylation can occur on the same sequence, (2) there is no overpropionylation, (3) N-terminal is always propionylated, (4) all lysines are modified (no underpropionylation), and (5) K37 us always unmodified (i.e., only propionylated). This constitutes the biologically relevant search space. The list of all theoretical variant×PTM combinations was generated using the *pyteomics* Python package (Levitsky et al., 2018).

**Figure 2.**
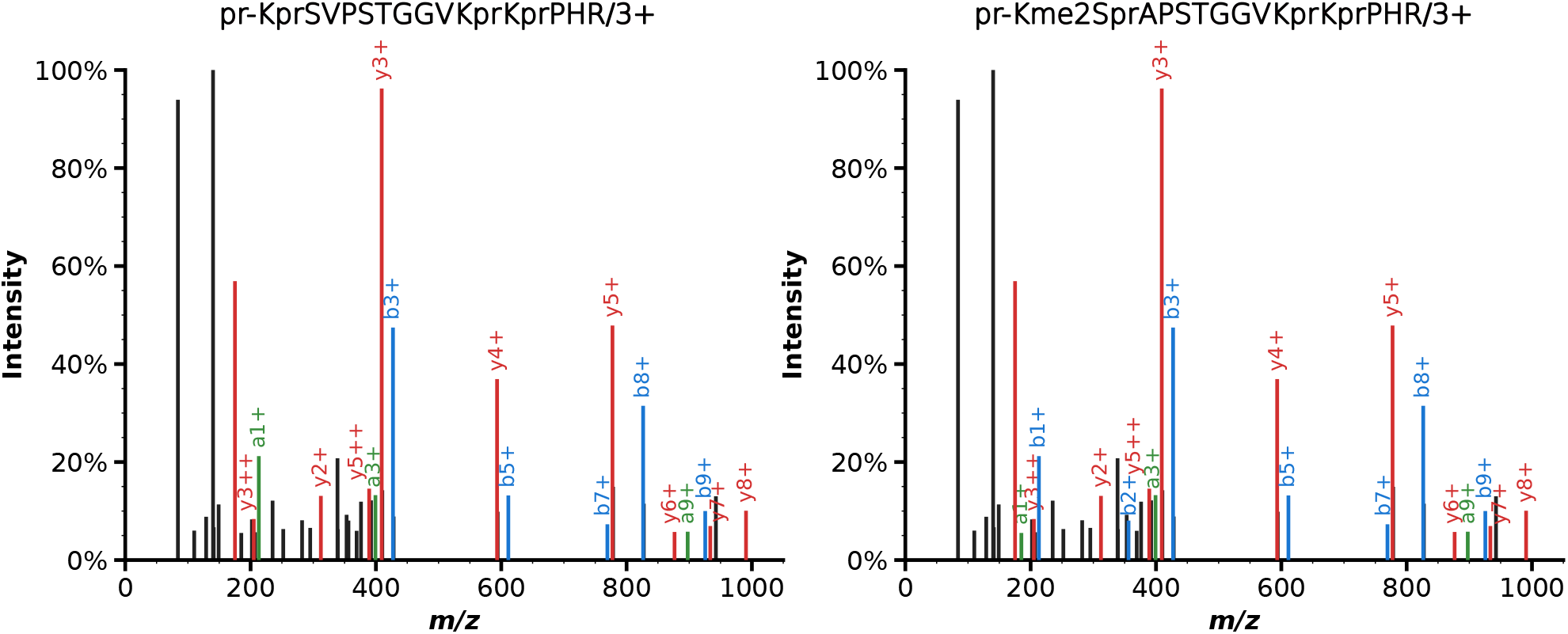
Overpropionylation makes histone MS/MS spectra interpretation ambiguous. The same MS/MS spectrum, first tentatively identified as unmodified H3mm13 K27-R40, was more confidently interpreted as H3.3 K27-R40 modified by K27me2/S28pr/K36pr/K37pr, with overpropionylation attributed to S28 when propionylation (S/T) was added to the list of allowable variable modifications.

**Figure 3.**
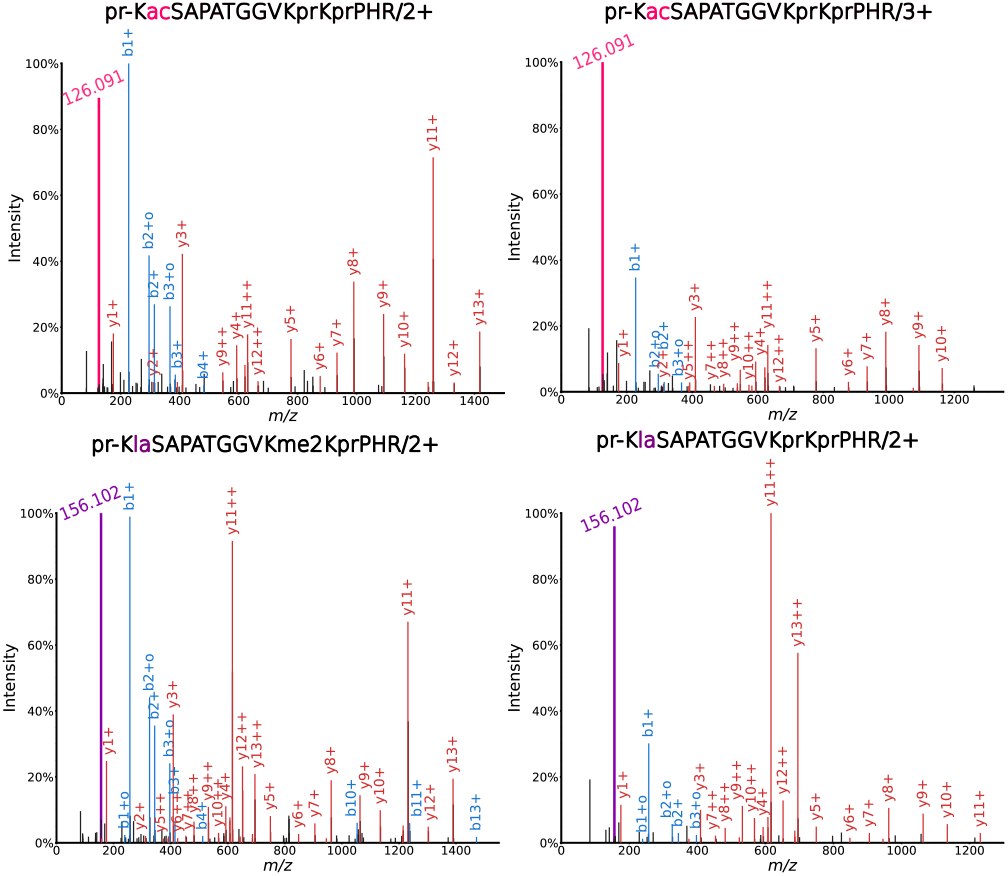
Cyclic immonium ion and b1 fragment ion are crucial in identifying H3K27acyl. The upper and lower panels show MS/MS spectra for H3cano K27ac-K36prK37pr and H3cano K27la-K36me2-K37pr, respectively, in 2+ (left) and 3+ (right) charge states. The CycIm ion corresponding to Kacyl is very intense when Kacyl is in the first position in the peptide. Acetylation and lactylation at H3K27 form intense characteristic CycIm ions at m/z 126.091 (pink) and 156.102 (purple), respectively. Fragment b1, which is stable when the peptide N-terminal is modified, is an important marker to validate acylation at H3K27, often representing the base peak. It is detected at very high intensity from peptides in 2+ forms (left spectra), but is less intense for peptides in 3+ forms (right spectra).

All DDA search results were filtered (rank 1 PSMs with a minimal score of 30 and minimal length of 6) and validated using Proline software (version 2.2) (Bouyssié et al., 2020). A spectral library was created in Skyline (MacLean et al., 2010) starting from four DAT files generated by Mascot searches of data acquired on a synthetic mixture containing variably modified H3K27-R40 and three technical replicates of the pooled biological samples spiked with the same synthetic mixture in DDA mode. A custom R script was used to swap a few spectra between the *redundant*.*blib* and *main*.*blib* library files to ensure that the reference spectra were taken from an ID very close to the apex. PRM files were imported into Skyline and every identification was manually checked for its predicted RT and *dotp* values. Only three fragment ions per identification were retained for quantification. Because distinct fragments were selected for variably modified peptides, it is only appropriate to compare the same PTM combination on H3 K27/K36 across samples.

Data obtained from Proline and Skyline were wrangled in R using *tidyverse* packages. Mirror, bar, and heatmap plots were generated using *ggplot2* and *patchwork* R packages.

### Data Availability

Mass spectrometry DDA and PRM data and their respective analysis files were deposited to the ProteomeXchange Consortium (http://www.proteomexchange.org) via Proteomics Identification Database (PRIDE) (Perez-Riverol et al., 2022) and Panorama Public (Sharma et al., 2018) partner repositories with the dataset identifiers PXD057347 and PXD057376, respectively. In addition, proteomics samples metadata were provided as Sample and Data Relationship File (SDRF) format (see also Supplementary Table S2) which have been generated using lesSDRF (Claeys et al., 2023) tool. The PRM analysis Skyline document is accessible on PanoramaWeb via https://panoramaweb.org/y11E4i.url. The scripts used to generate the data and figures are available under https://github.com/HijaziHassan/VariantPaper.

## RESULTS AND DISCUSSION

### Mascot tentatively identified the mouse-specific variants H3mm7 and H3mm13 in histone samples extracted from mouse brain

Several mouse-specific variants were identified as incorporated into chromatin (Maehara et al., 2015). Among them, the transcripts of H3mm7 and H3mm13 appeared to be very abundant in mouse brain and testis. We then investigated whether the corresponding proteins could be identified in our histone samples processed by in vitro propionylation. H3mm7 and H3mm13 differ from H3.3 by one or two amino acids after Lys27 (see sequences in Table 1). When analyzing histone H3 from mouse brain, we tentatively identified these two variants in several modified forms of the stretch K27R40, all with a Mascot score over 30 (Table 1). A universal spectrum identifier (USI) (Deutsch et al., 2021) for each of these tentative identifications is provided in Supplementary Table S3. In all cases, most of the higher-intensity experimental fragments could be interpreted in terms of y and a/b fragments, which would a priori support these peptide identifications. However,the proteomic analysis of histones submitted to chemical propionylation has specificities, and a closer look at MS/MS spectra may be required.

**Table 1.**
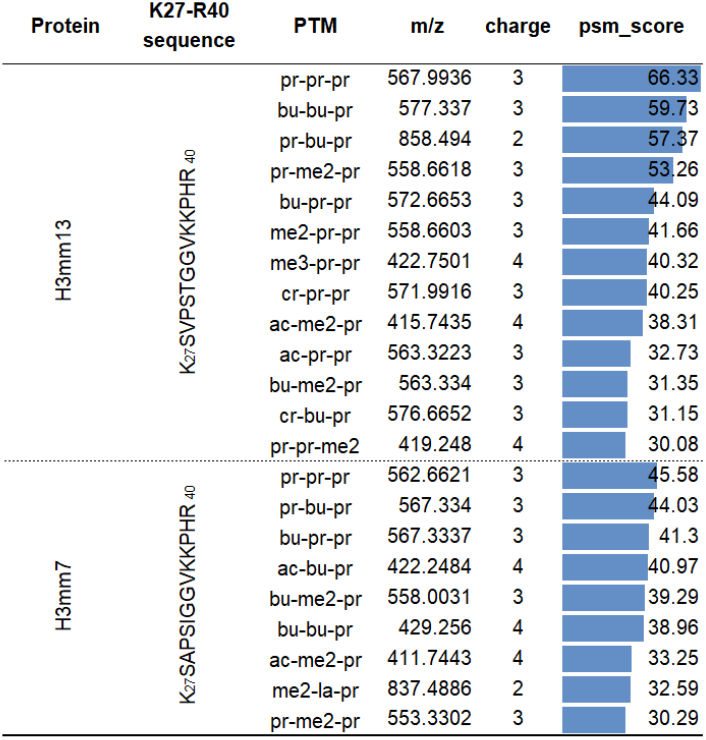
Variably modified peptides from mouse-specific H3 variants H3mm7 and H3mm13 tentatively identified in mouse brain with a Mascot score over 30.

### The number of isobaric H3K27-R40 peptidoforms increases dramatically when multiple acylations, sequence variants and overpropionylation are allowed

In classical proteomic analyses, the precise mass measured for a peptide usually efficiently restricts the list of potential matching theoretical sequences (Elhamraoui et al., 2024). To precisely evaluate how well this rule applies when analyzing histones, we calculated the number of isobaric H3 K27-R40 peptidoforms resulting from combinations of PTMs, sequence variations, and uncontrolled propionylation. First, we generated a list of all the possible peptidoforms when considering Nterminal and lysine propionylation as variable modifications alongside classical endogenous PTMs (Kac and Kme1/2/3) on the three lysines in the K27-R40 stretch from canonical H3 and variant H3.3 only (Figure 1a; red bars). This list delimits the whole search space explored by an interpretation program, and includes completely unmodified peptides. The list was then extended by adding three novel acylations: lactylation, crotonylation, and hydroxybutyrylation (+Kla/cr/hb) (Figure 1a; blue bars). Next, more masses were appended by including mouse-specific histone variants (+H3mm7/13 variants) (Figure 1a; green bars). Finally, overpropionylation of S/T was factored in, which considerably increased the number of peptides with the same exact mass (Figure 1a; same color code but with translucency, and Supplementary Table S4). All the theoretical masses that matched high numbers of variant×PTM combinations emerged after considering overpropionylation.

As a showcase, we followed the stepwise increment of the number of variant×PTM combinations with a mass of 1684.96244 amu (Figure 1b). Only three variant×PTM combinations strictly match this mass when classical PTMs are considered. This number increases to 15 combinations after including novel lysine acylations (+Kla/cr/hb); to 49 combinations when mouse-specific variants (+H3mm7/13) are included; and to 306 combinations when overpropionylation on S/T (+Overprop) is considered. This theoretical calculation illustrates how overpropionylation can easily lead to an exceedingly large number of isobaric peptides, complexifying data interpretation and increasing the chances of producing false positive identifications.

Finally, based on prior knowledge, we narrowed the search space by only considering the variant×PTM combinations of H3 K27-R40 peptides that we often deliberately seek. Thus, only combinations that contained no more than one novel acylation – among lactyl, crotonyl and hydroxybutyryl – were retained, because these PTMs are generally of lower abundance. We also kept sequences that were endogenously unmodified at K37 (i.e., propionylated), because this residue has rarely been reported as modified (Vai et al., 2024). Finally, we kept peptides that were not over- or underpropionylated (i.e., all lysines were modified but no other residues). The comparison of Figures 1a and Figure 1c shows how the maximum number of variant×PTM combinations matching the same peptide mass substantially drops when applying these constraints. This theoretical assessment illustrates that biologically relevant combinations need to be carefully extracted from a very specific search space when dealing with multiple lysine PTMs, sequences containing more than one such residue, exhibiting subtle amino acid variations and prone to overpropionylation.

To demonstrate the detrimental effect of S/T overpropionylation on histone peptide identification at an experimental level, we analyzed samples of histone H3 extracted from mouse brains. Following data interpretation with Mascot against a histone sequence database containing mouse-specific variants (El Kennani et al., 2017), the program tentatively identified peptide K27-R40 from H3mm7 and H3mm13 with variable PTMs on K27/K36/K37. One example (Figure 2) corresponds to an MS/MS spectrum assigned to sequence K_27_prSVPSTGGVKprKprPHR_40_ from H3mm13, which is the endogenously unmodified mouse-specific histone variant. It is tempting to validate this identification, given that all b and y fragments of masses above m/z 400 are labeled. However, although b3 is detected, the N-terminal Kpr can only be confirmed by the detection of an a1 fragment. Interestingly, when allowing S/T propionylation during interpretation of this dataset, a higher-score identification (score 79 versus 58) was obtained for K_27_me2SprAPSTGGVKprKprPHR_40_ from H3.3. In this case, fragments b1, b2, and b3 are labeled in the experimental spectrum, in strong support of the dimethylation on the first lysine and the propionylation on the serine in second position (Figure 2). This propionylation of the S residue thus led to erroneous identification of H3mm13, as the mass difference between A and V residues is identical to the shift induced by dimethylation (+28.0313 Da) at H3.3K27. This side reaction can be efficiently eliminated by treating peptide samples with hydroxylamine at pH 12 Meert et al. (2016); **?**. For the results hereafter on endogenous histone samples, we systematically implemented this reversepropionylation step in our histone processing protocol before MS analysis.

In conclusion, due to numerous cases of isobaric amino acid variant×PTM combinations, the precisely measured mass may often be a weak filter when seeking to identify modified histone peptides prepared by in vitro propionylation and trypsin digestion, especially when additional novel lysine acylations and histone variants are considered.

### Fragment b1 and cyclic immonium ions are crucial to confirm the identity of PTMs on H3K27

In the fragmentation spectrum of a tryptic peptide, unless the peptide N-terminus is modified, the b1 fragment is normally not observed due to chemical instability (Michalski et al., 2012). Following in vitro propionylation of the N-termini of histone peptides, we expect to systematically observe the b1 fragment in MS/MS spectra (Figure 3). In addition, a modified lysine in the fragmented peptide produces an immonium and an immonium-related ion that has lost NH3 (i.e., CycIm ion) (Zolg et al., 2018; Hseiky et al., 2021; Wan et al., 2022). For example, acetylated and lactylated lysines produce characteristic CycIm ions at 126.091 and 156.102 m/z, respectively. The closer the acylated lysine is to the Nterminus, the more easily the CycIm ion is formed and the greater its intensity in the MS/MS spectrum (Madden et al., 1991; Couttas et al., 2008; Muroski et al., 2021). Interestingly, due to the frequent occurrence of the RK stretch, lysine is often at the first position of the sequence in histone peptides.

We scrutinized the MS/MS fragmentation spectra of endogenous and synthetic peptides corresponding to tens of variant×PTM combinations covering H3 K27R40 with various acylations at K27 and no modification or dimethylation at K36. We detected b1 in all MS/MS spectra, although at varying intensities depending on the charge state of the precursor ion. The b1 fragment ion was very intense in MS/MS fragmentation spectra from peptides in the 2+ charge state, but much less intense with peptides in 3+ and 4+ forms, as is shown in Figure 3 for two canonical H3 K27-R40 peptidoforms: pr-K_27_acSAPATGGVKprKprPHR_40_ and prK_27_laSAPATGGVKme2KprPHR_40_ in 2+ and 3+ charge states (also see Supplementary Table S5 containing USIs of spectra for other variant×PTM combinations). In addition, the CycIm ion produced by H3K27acyl was either the base peak or a runner-up (Figure 3), indicating the importance of this ion in revealing the identification and localization of a PTM at the first lysine of the peptide sequence.

Overall, the b1 fragment and CycIm ions provide critical information in an MS/MS spectrum making it possible to confirm the identity of H3K27acyl and to flag false positives in the case of isobaric peptides.

### Preprocessing options applied to MS/MS spectra may eliminate the low-mass range

Because b1 fragments and CycIm ions are central to confirming histone peptide identification, the information they carry must be included in the MS/MS spectra during raw data processing. We routinely run database searches with the Mascot program, which is recommended for histone PTM identification (Yuan et al., 2014), after preprocessing RAW files with our in-house developed program MGFBoost or Mascot Distiller. In the latter tool, parameters that are usually not reported include how peak lists are extracted from the raw data in formats such as MGF (Mascot Generic Format). These parameters include “intensity values: area or S/N”, which influences the relative intensity of lowand high-mass fragments, and “output MS/MS fragment as: MH+ or m/z”, which can deconvolute multiply-charged fragments into their singly-charged counterparts. We analyzed synthetic peptides spanning residues K27-R40 from H3 variants carrying variable modifications, and compared the relative intensities of their fragment ions with each combination of the above Mascot Distiller parameters. Two peptidoforms (pr-K_27_acSAPATGGVKprKprPHR_40_ [2+] and pr-K_27_acSAPATGGVKme2KprPHR_40_ [3+]) of canonical H3 histone were used to illustrate the impact of these parameters during processing for peaklist generation. The processed spectra are shown alongside the raw spectra to aid visual comparison (Figure 4). Intense low-mass ions were more pronounced when the S/N option was selected regardless of the chosen “output fragment as” option (Figure 4; SN_MH and SN_moz). In contrast, the “Area” option almost completely eliminated this low-mass region (Figure 4; Area_MH and Area_moz). Choosing to de-charge the fragment ions (”MH” option) led to aggregation of the signals of 2+ fragment ions in their 1+ form in the case of triply charged pr-K_27_acSAPATGGVKme2KprPHR_40_ peptide. To maintain control over the preprocessing steps, we use our in-house tool, MGFBoost, before running Mascot database searches. MGFBoost performs multiple tasks (detailed elsewhere; paper in preparation), and most importantly for histones, it retains all the features of the raw spectrum (Figure **??**; MGFBoost). When used in conjunction with the pClean R package (Deng et al., 2019) for fragment ion deconvolution and noise reduction, the spectra are very comparable to those obtained when applying the “Area_MH” option in Mascot Distiller, except that the intense ions in the low-mass region are retained (Figure **??**; MGF-Boost_pClean). We also compared results obtained with other software tools widely used in the MS community, namely MaxQuant (Andromeda; v2.4) (Cox et al., 2011) and FragPipe (MSFragger; v21.1) (Kong et al., 2017). These results are presented in Supplementary Data Section 2 and Figure S1 which shows that these two programs retain the low-mass range of MS/MS spectra. In summary, the relative intensities of the fragment ions can vary depending on the parameters chosen when converting a RAW file into a peak list. Using our custom tool MGFBoost (with or without pClean before Mascot data interpretation), Fragpipe, or MaxQuant allows for faster manual validation of modified histone spectra as they retain b1 and CycIm ions. We recommend taking care to retain the low-mass region containing these ions that are critical for correct identification of histone peptides and their lysine PTMs.

**Figure 4.**
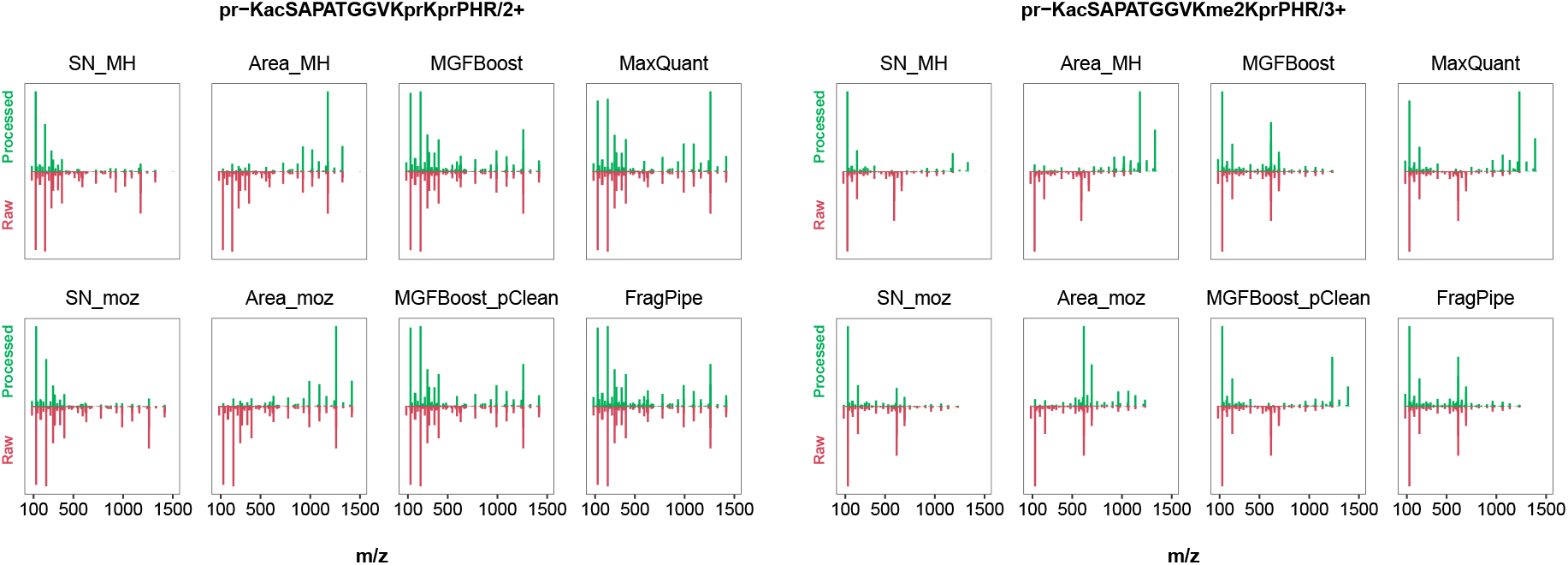
Preprocessing parameters can drastically alter the MS/MS spectra submitted for peptide identification. Data for two peptidoforms with different charge states from canonical H3: K27ac-K36pr-K37pr/2+ and K27ac-K36me2-K37pr/3+ are shown. The first four panels represent the four combinations of Mascot Distiller processing options. The 5th and the 6th panels represent MGFBoost with and without the pClean preprocessing option.

### Retention time behavior of variably modified H3 K27-R40 stretches

Our previous observations on the production of intense discriminant b1 and CycIm ions are valid in the case of MS/MS spectra acquired for a single modified histone peptide. However, co-elution and thus co-fragmentation of histone peptides with similar m/z ratios occur quite frequently, leading to composite MS/MS spectra. The RT constitutes an orthogonal piece of information that can be used to discriminate histone peptides (Yuan et al., 2015, 2018). We therefore determined the relative chromatographic behavior of 40 variant×PTM combinations for the K27-R40 stretch from the four H3 sequences, and monitored the variant elution order relative to canonical H3 peptides (see Supplementary Table S1). The combinations studied included variable modifications on K27 (ac and me1/2/3), with either me2 or pr on K36, and pr on K37. We observed that the RTs remained unchanged across the three variants relative to the canonical H3 RT, regardless of the PTM combination, with median differences of -0.64, 3.68, and 8.65 min for H3.3, H3mm13, and H3mm7, respectively (Figure 5a). From the constant relative RTs observed in Figure 5a, the relative LC behavior of other sequence×PTM combinations can be established. For instance, the RT difference between H3 K27ac-K36prK37pr and H3 K27ac-K36me2-K37pr peptides was approximately 7.2 min, due to the replacement of dimethylation with propionylation at K36. This consistency can be used to confidently anticipate the RT shift for H3 K27la-K36me2-K37pr following detection of H3 K27laK36pr-K37pr, thus eliminating the need to produce multiple synthetic reference peptides. The RT therefore provides an additional filter to ensure correct peptide identifications.

**Figure 5.**
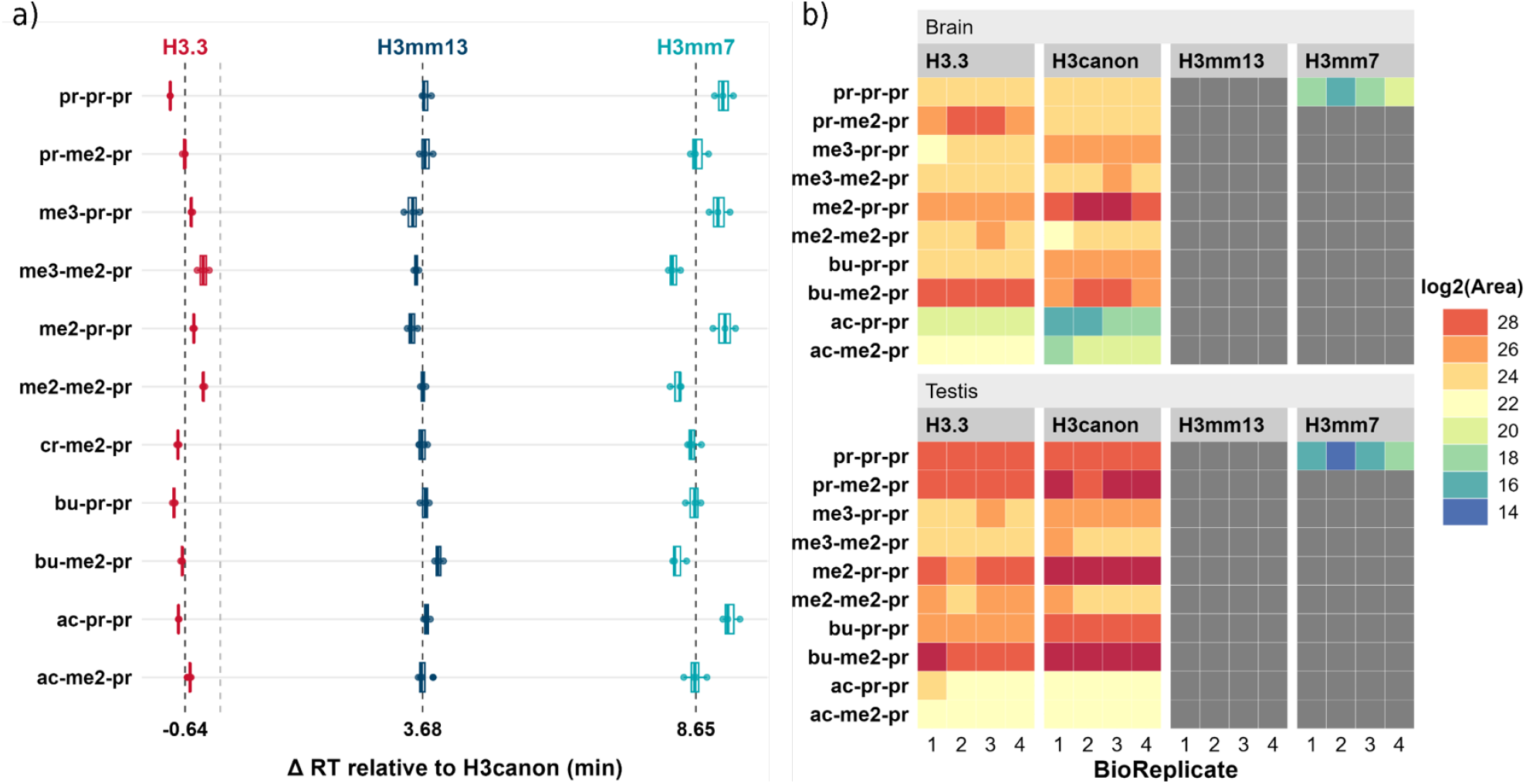
Elution trend for 40 H3 K27-R40 variant×PTM combinations and their targeted analysis in mouse brain and testis. (**a**) Retention time differences for various H3 K27-R40 variant×PTM combinations relative to their H3canonical counterparts. The vertical light grey dotted line represents the reference RT (0 min) corresponding to H3canonical K27-R40; the black dotted lines represent the median relative ^Δ^RT calculated from three or four technical replicates. (**b**) Heatmap showing the MS/MS-based abundance of H3 K27-R40 variant×PTM combinations targeted by PRM in histone samples extracted from mouse brain and testis. The log2-transformed intensity values of 3 fragment ions per peptidoform were summed and normalized relative to the total ion current (TIC) area. Undetected peptides are indicated in dark gray.

### PRM and middle-down analyses fail to convincingly validate the presence of H3mm7 and H3mm13 histone variants

The above comprehensive analyses laid the groundwork for robust MS analysis of H3 K27-R40 peptidoforms in histones extracted from mouse brains and testes. We targeted variant×PTM combinations with classical PTMs at K27 (ac/me1/2/3 in addition to pr) with either pr or me2 at K36, but leaving K37 propionylated. This subset of 10 combinations was chosen based on our observations of canonical H3 and H3.3 and their high abundance reported in the literature (Tvardovskiy et al., 2017). An isolation list containing the m/z of the most abundant charge state of each peptidoform and their respective RTs was defined and used in targeted analyses in PRM mode (See Supplementary Table S1). We generated a curated spectral library from DDA data acquired on pooled histones extracted from testis and brain tissues spiked with a mixture of synthetic peptides derived from the different H3mm7 and H3mm13 K27-R40 peptidoforms targeted. Thus, high-quality reference MS/MS fragmentation spectra were available for those mouse-specific novel histone peptides. In addition, both the pooled sample and the four biological replicates prepared for the targeted experiment were spiked with iRT peptides. Signals from these peptides were used to build an iRT calculator in Skyline, to allow accurate prediction of RTs despite any LC variations. These dimensionless RTs determined for the target peptides facilitate lab-to-lab transferability of this method.

In the seminal paper describing their discovery, high levels of H3mm7 and H3mm13 transcripts were detected in the brain and testes of 8-week-old mice using the 3 ^*′*^seq method (Maehara et al., 2015). Nonetheless, none of the PTM combinations of H3mm13 and H3mm7 targeted in this study, apart from the unmodified form of H3mm7 K27-R40 (i.e., K27pr-K36pr-K37pr) were more than weakly detectable at the MS/MS level, and not at all at the MS1 level Figure 5b. Unexpectedly, H3mm13 peptidoforms were never detected. This H3.3 subvariant, if it exists at the protein level, may thus be present at less than the lower limit of detection in the testis and brain in 3-month-old mice. These observations from the targeted bottom-up analysis were further supported by a middle-down analysis of histones extracted from mouse testis using the ETD fragmentation mode. Following Glu-C digestion, large peptides covering the 50amino-acid-long H3 N-terminus were analyzed. The canonical H3, H3.3, and testis-specific H3 (H3t) were all unambiguously identified in MS/MS spectra based on discriminating *c* and *z* fragments produced during ETD. Importantly, whereas our former targeted analysis omitted variant H3t, which is only expected in the testis, this exploratory middle-down analysis identified it based on several variably modified peptides (Supplementary Data Section 3). The sets of PTMs observed on K27/K36 of variants H3.3 and H3t further confirmed the relevance of the PTM combinations selected for our former PRM analyses. Certain ambiguous spectra lacking fragments between K27 and K36 led to tied scores for H3mm7 and canonical H3 on the one hand and H3mm13 and H3.3 on the other hand, but we were unable to validate any spectrum clearly identifying H3mm13 or H3mm7. Taken together, our data suggest that *H3mm13* is likely a pseudogene, whereas H3mm7 is barely detectable even with the enhanced sensitivity of targeted MS methods.

## CONCLUSIONS

Histones exist in several sequence variants and bear an exquisitely complex array of PTM combinations mainly localized on their multiple lysine residues. The H3K27-R40 segment alone contains three lysine residues and slight amino acid substitutions that result in numerous isobaric variant×PTM combinations. Our investigation of this combinatorial space revealed that in addition to endogenous modifications and sequence variants, uncontrolled propionylation at S/T residues significantly increases the number of isobaric variant×PTM combinations. Consequently, care must be taken when reporting histone PTM identifications. First, if overpropionylation is detected, it should absolutely be removed by a final treatment step with hydroxylamine (Meert et al., 2016). In addition, the b1 fragment and CycIm ion in MS/MS spectra are highly significant for the confirmation of the presence of the acyl modification at H3K27, and more generally at lysines in the first position of a histone peptide. We observed that CycIm ions were consistently intense regardless of the precursor ion charge state. However, this intensity can be reduced during the conversion of raw data to peak lists, depending on the software used. Therefore, it is critical to control the preprocessing step to ensure they remain present in the final interpreted MS/MS spectra. In addition, it would be relevant to more systematically consider their detection when scoring matches. This is already possible when using Andromeda (Zolg et al., 2018). To complete the criteria useful for validating the identification of histone peptides, the RT behavior of variably modified K27-R40 stretches corresponding to four H3 variants was established. We used this combined information to perform targeted PRM analyses on histones from mouse brain and testis, revealing only very weak detection of the unmodified H3mm7 K27-R40 stretch and no detection of H3mm13 at the protein level. This study is the first LC-MS/MS analysis targeting different peptidoforms of these mouse-specific variants using bottom-up and middle-down approaches. It may be interesting to explore whether these reported mouse-specific variants exist at the protein level in specific conditions, such as in aged mice, in mouse mutants, or in specific cell types.

## Notes

### Competing Interest Statement

The authors have declared no competing interest.

### Summary of Updates

We have removed unneccessary filters used to generate the data used to produce Figure 1. We update it and this reinforces our messages.

