## Supplementary Data for "Mind Your Spectra: Points to be Aware of when Validating the Identification of Isobaric Histone Peptidoforms"

#### Section 1. Animals used for this research

##### Testis

For testis collection, adult C57BL/6 male mice (2.5- to 3-month-old) were used and all procedures were subjected to local ethical review (Comité d'Ethique pour l'Experimentation Animale, Université Paris Descartes, registration number CEEA34.JC.114.12, APAFIS 14214-2017072510448522v26 to Julie Cocquet, Université Paris Cité, INSERM, CNRS, Institut Cochin, F-75014 Paris, France).

##### Brain

Brain tissues were obtained from 2.5- to 3-month-old male mice maintained on C57BL/6J background at LNCA. Animal studies were conducted in accordance with French regulations (EU Directive 2010/63/UE–French Act Rural Code R 214-87 to 126). The animal facility was approved by veterinary inspectors (authorization no. E6748213) and complies with the Standards for Human Care and Use of Laboratory Animals of the Office of Laboratory Animal Welfare. All procedures were approved by local ethics committee (CREMEAS) and French Research Ministry (no. APAFIS#504-2015042011568820_v3). Mice were housed in a controlled-temperature room maintained on a 12 h light/dark cycle. Food and water were available ad libitum. Mice were killed by cervical dislocation and brain hemispheres were rapidly dissected, snap frozen and stored at −80 °C.

#### Section 2. How do MaxQuant and Fragpipe preprocess MS/MS mass spectra?

For the sake of generalizability, we also included in the comparison of Mascot preprocessing parameters and MGFBoost other software tools widely used in the MS community, namely MaxQuant version 2.4 (Andromeda)[1] and FragPipe version 21.1 (MSFragger)[2]. MaxQuant deconvolutes multiply charged fragment ions into their singly protonated form but unlike Mascot Distiller, it maintains the low mass fragment ions (Figure S1; MaxQuant). This behavior is coherent with the fact that CyIm ions contribute to the final Andromeda score. MSFragger in FragPipe allows the user to output an .mzML peak list suffixed with either "_calibrated" or "_uncalibrated" depending on whether the mass calibration parameter is selected or not. We chose the latter as it is the default output and it is automatically associated with the visualization viewer (FragPipe-PDV)[3] found in the FragPipe suite in case both files are present. The mirror plots revealed almost the same output as MGFBoost, indicating minimal changes, if any, during the processing step from raw to peak list transformation (Figure S1; FragPipe). However, it should be noted that the "_calibrated.mzML" file contains deconvoluted fragment ions and shows a fragmentation profile similar to MaxQuant and MGFBoost_pClean processed spectra.

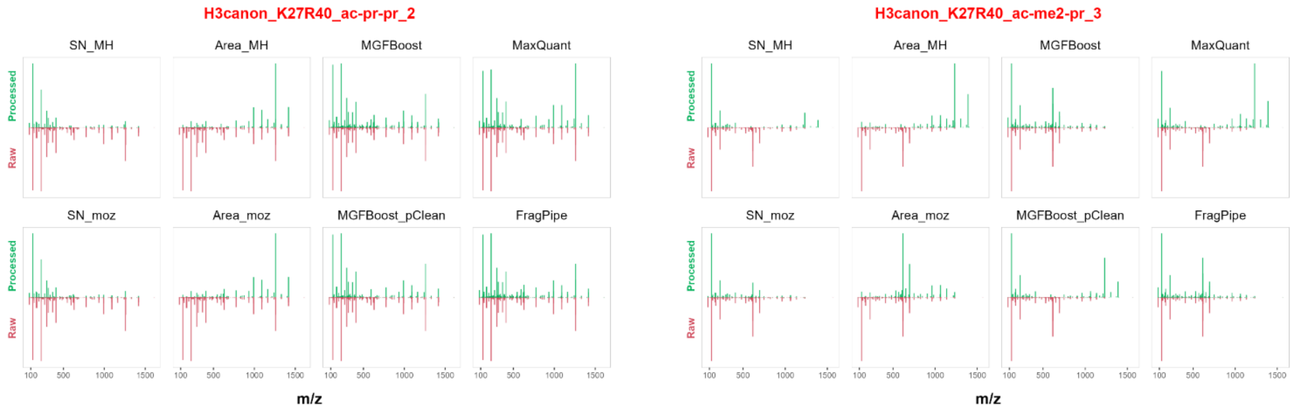


Figure S1. **Preprocessing drastically impacts the shape of MS/MS spectra submitted to peptide identification**. The first four panels represent the four combinations of Mascot Distiller processing options. The 5th and the 6th panels represent MGFBoost with and without pClean preprocessing options. The last two panels correspond to the processing behavior of Andromeda and MSFragger that are embedded in MaxQuant and FragPipe software, respectively. The peak list files were recovered after a DDA search and converted to *mgf* format. For Andromeda (MaxQuant v2.5), the *.apl* file containing the peak list was converted to *mgf* using MGFBoost, whereas the *mzML* file generated from MSFragger (Fragpipe v21.1) was converted using MSConvert (v3.02).

In summary, the relative intensities of the fragment ions can vary depending on the settings chosen when converting a RAW file into a peak list. Using either our custom tool MGF-Boost with or without pClean before Mascot data interpretation, FragPipe, or MaxQuant allows for faster manual validation of modified histone spectra as they retain CycIm ions. In histone PTM analysis, we recommend taking care to retain the low-mass region containing characteristic CycIm ions that are critical for accurate identification and/or localization of lysine PTMs. Interestingly, FragPipe also allows for the discovery of novel diagnostic feature especially for more complex and labile PTMs such as glycans that produce different characteristic ions[4].

### Section 3. Middle-down Analyses of canonical H3, variants H3.3 and TSH3

#### N-term Sequences of canonical H3 and its variants.

| | |
|---|---|
| **H3.1** | ARTKQTARKSTGGKAPRKQLATKAARKSAPATGGVKKPHRYRPGTVALRE |
| **H3.3** | ARTKQTARKSTGGKAPRKQLATKAARKSAPSTGGVKKPHRYRPGTVALRE |
| **TS H3** | ARTKQTARKSTGGKAPRKQLATKVARKSAPATGGVKKPHRYHPGTVALRE |
| **H3mm7** | ARTKQTARKSTGGKAPRKQLATKAARKSAPSIGGVKKPHRYRPGTVALRE |
| **H3mm13** | ARTKQTARKSTGGKAPRKQLATKAARKSVPSTGGVKKPHRYRPGTVALRE |

#### Ambiguous spectra with the same score between H3mm7/H3.1 and H3mm13/H3.3

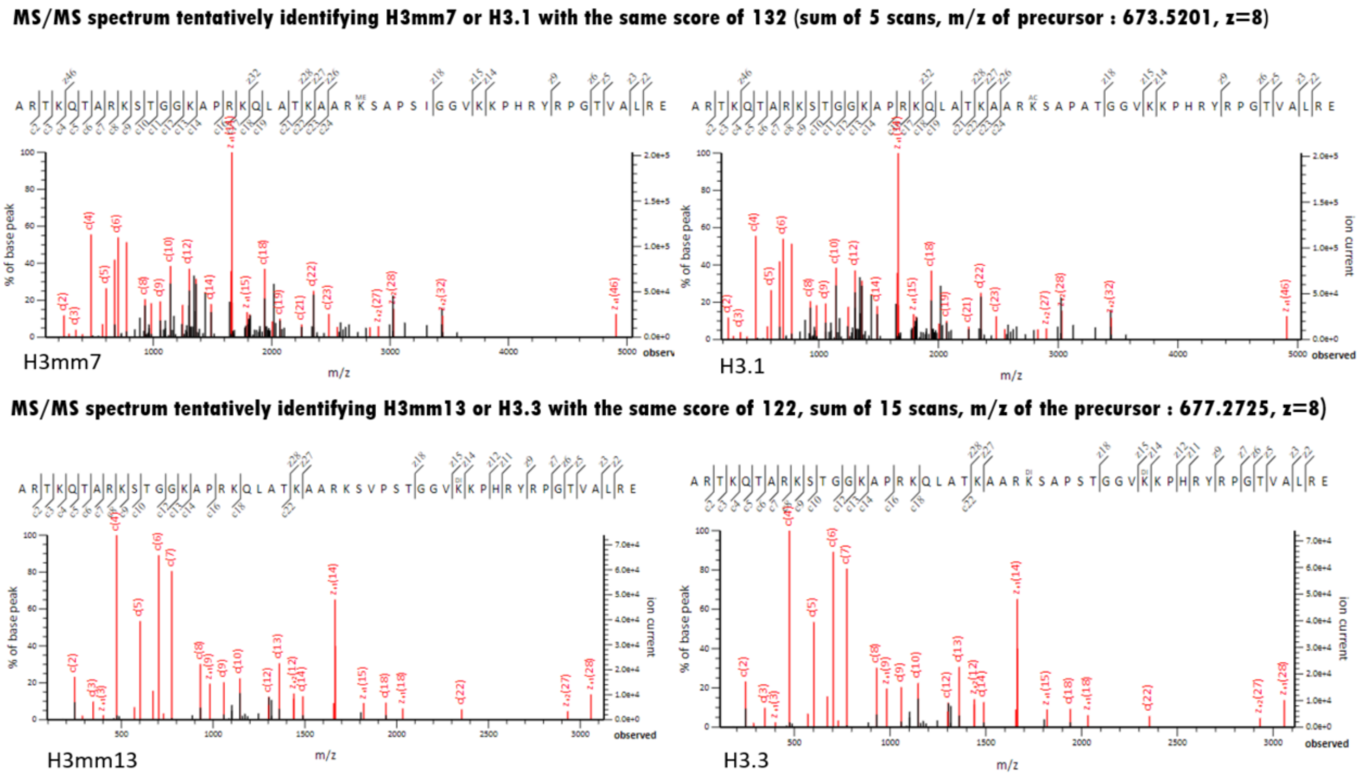


**Figure S2. Ambiguous spectra with the same score between H3mm7/H3.1 and H3mm13/H3.3.** The identification of H3mm7 and H3mm13 is not validated because there is a lack of c/z fragments surrounding the residues that differentiate these variants from H3.1 and H3.3.

#### Selected Identified MS/MS Spectra

### a) H3.1

*K27me2-K36me2-K37un*

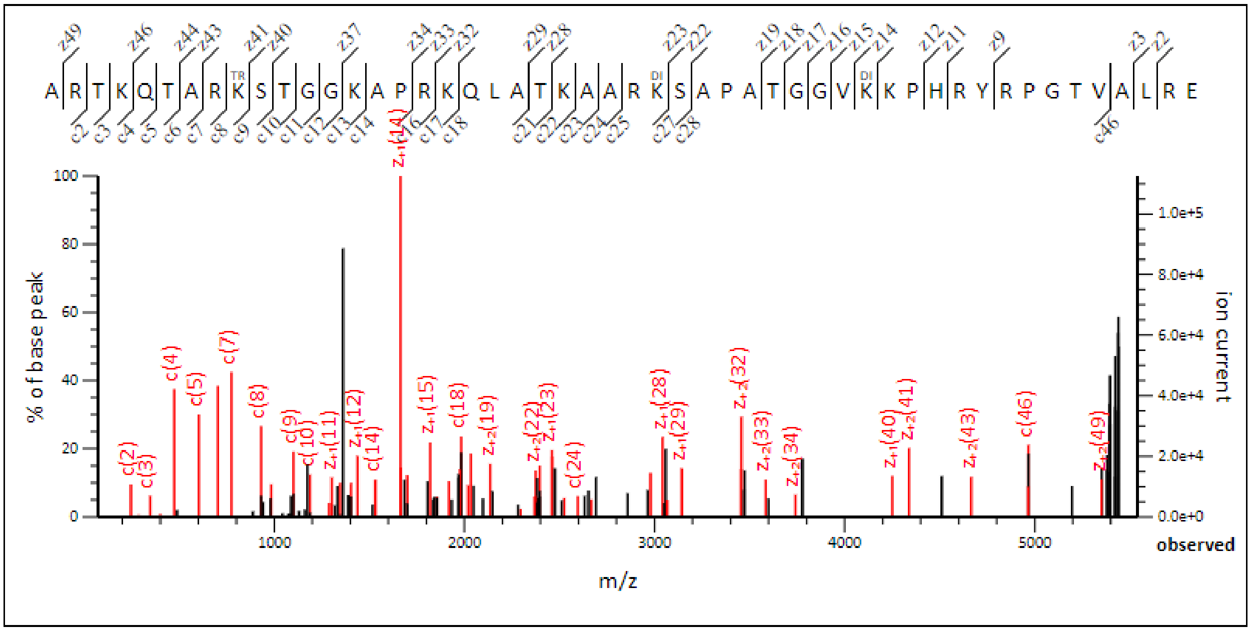


##### b) TSH3

*b.1. K27un-K36un-K37un*

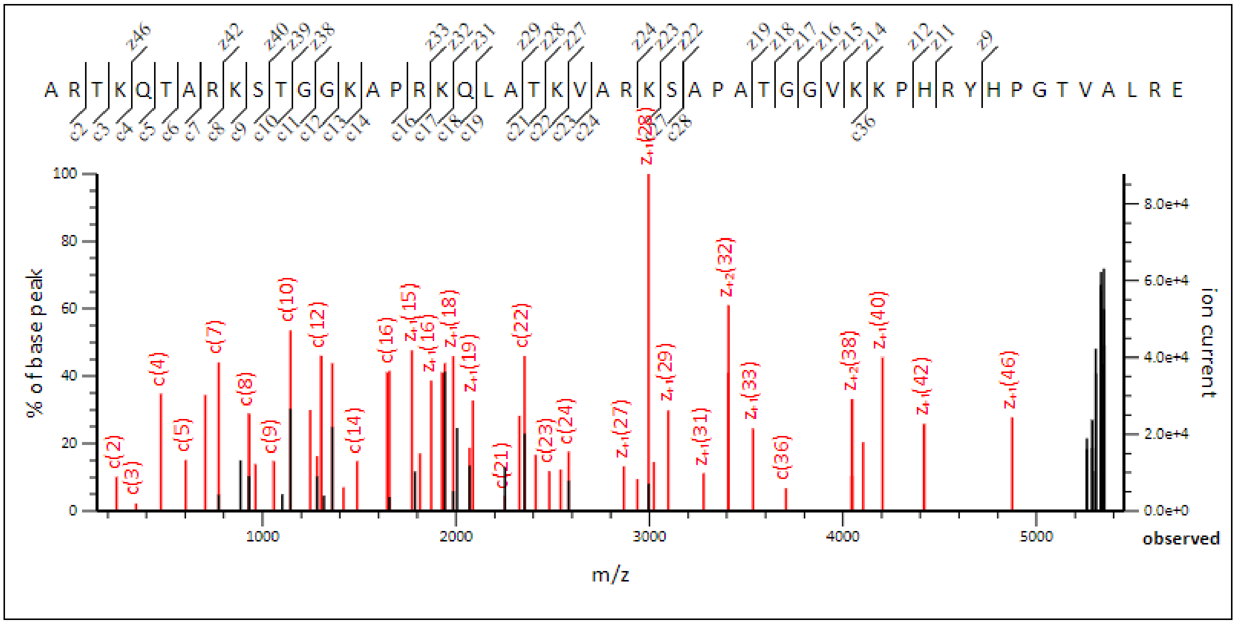

*b.2. K27me1-K36me1-K37un*

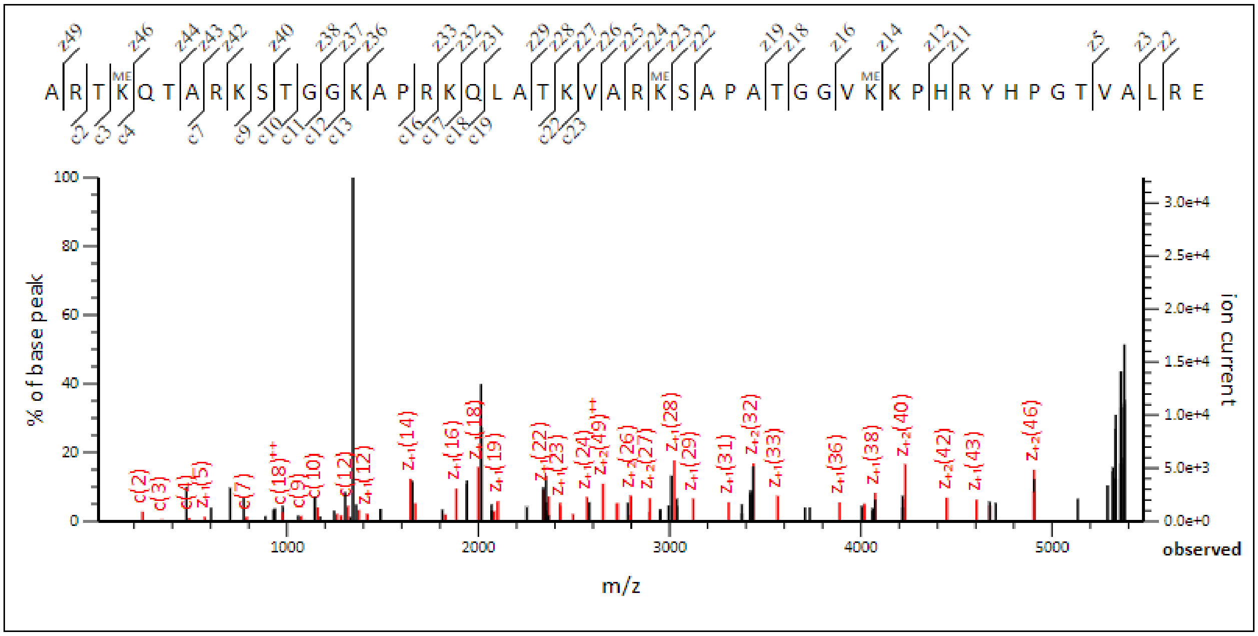


*b.3. K27me2-K36me2-K37un*

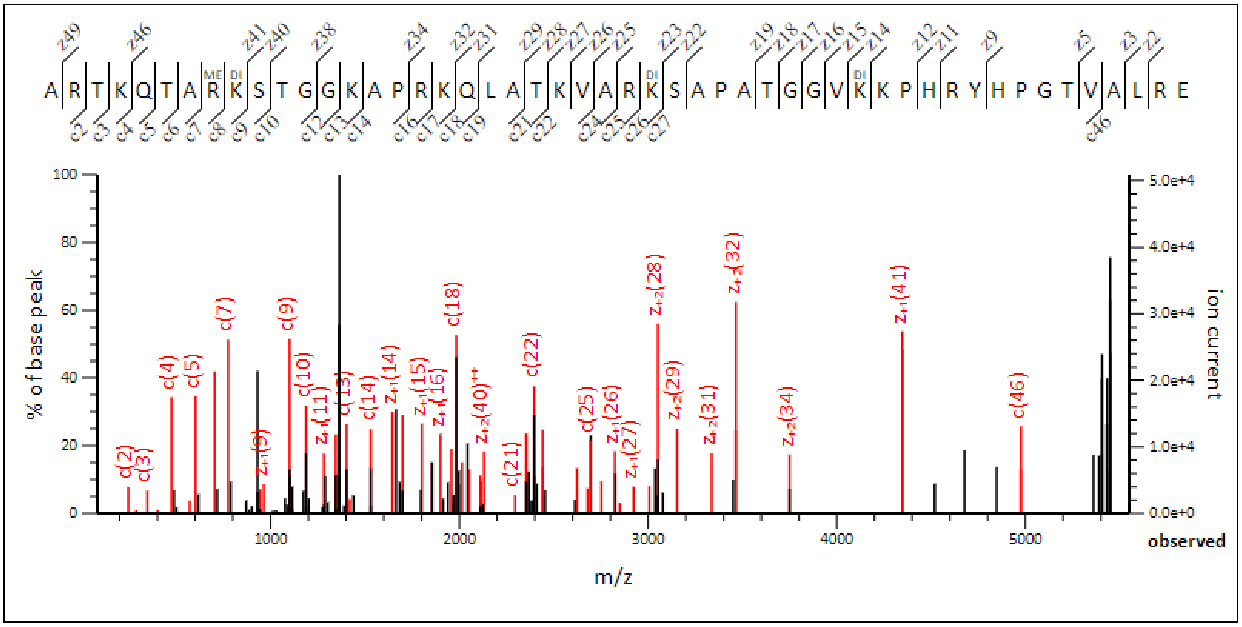

*b.4. K27me3-K36un-K37un*

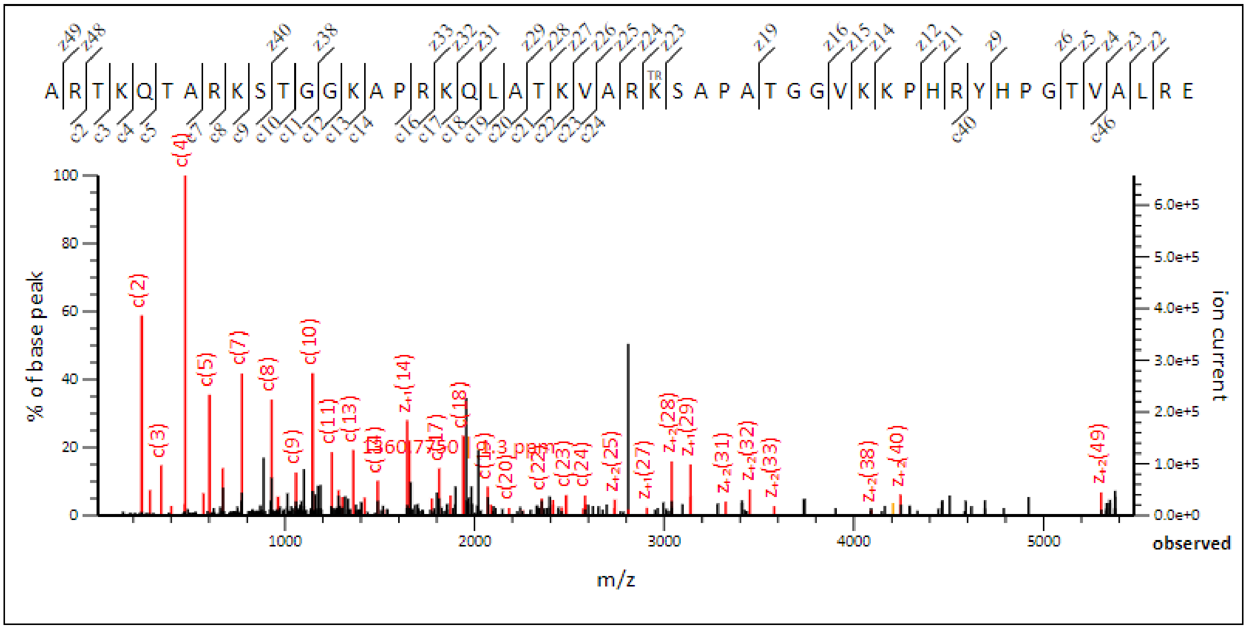


## c) H3.3

*c.1. K27un-K36un-K37un*

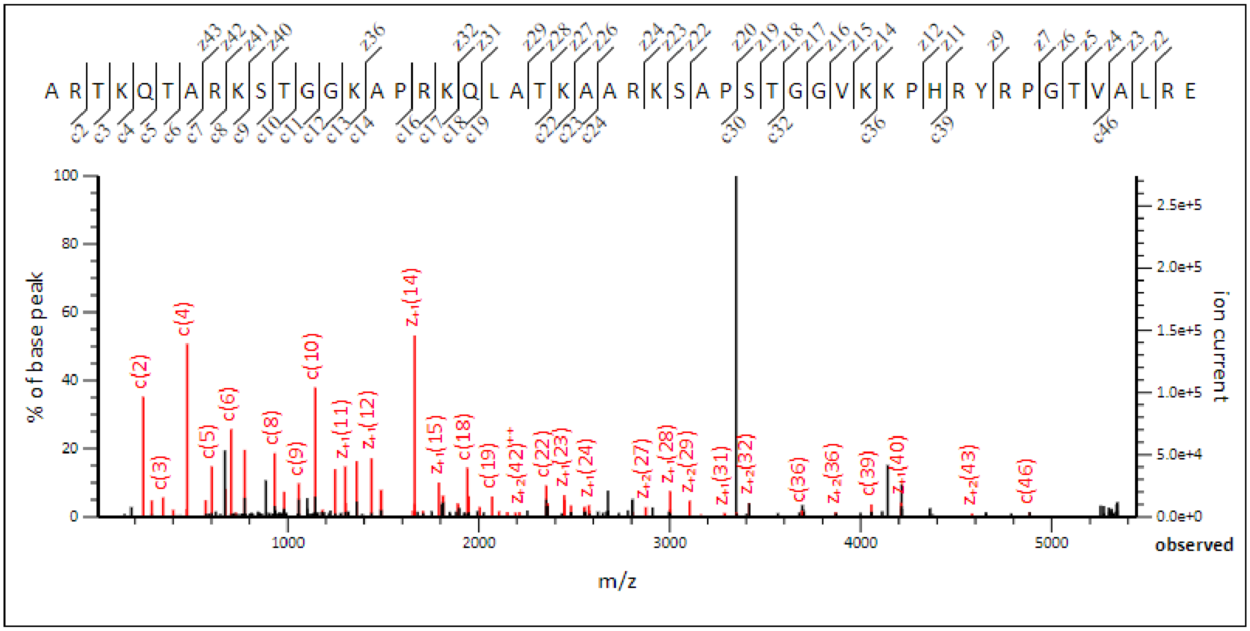

*c.2.K27me3-K36un-K37un*

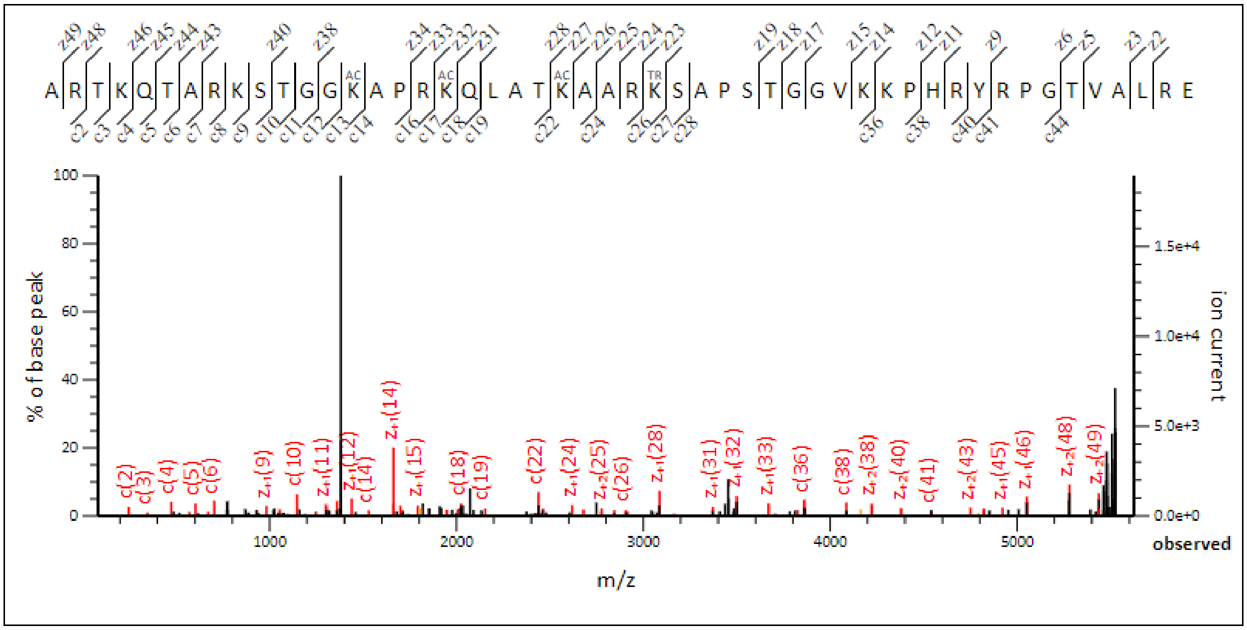


Figure S3. **Selected Identified MS/MS Spectra acquired by middle-down analysis.** (a) H3.1, (b) testis-specific H3 (TSH3) and (c) H3.3 were well identified by several PTM combinations covering residues 1-50. Only PTMs present on K27, K36 and K37 are indicated in the subtitles.
